# *TIC236* gain-of-function mutations unveil the link between plastid division and plastid protein import

**DOI:** 10.1101/2021.04.25.441383

**Authors:** Jun Fang, Bingqi Li, Lih-Jen Chen, Vivek Dogra, Shengji Luo, Pengcheng Wang, Inhwan Hwang, Hsou-min Li, Chanhong Kim

**Author notes:** These authors contributed equally to this work (Jun Fang, Bingqi Li).

## Abstract

The chloroplast translocons TOC75 and TIC236 are homologs of the bacterial translocation and assembly module (Tam) A and TamB involved in protein export. Here, we unveil a TIC236-allied component, the chloroplast outer membrane protein CRUMPLED LEAF (CRL), absence of which impairs plastid division and induces autoimmune responses in *Arabidopsis thaliana*. A forward genetic screen aimed at finding *crl* suppressors revealed multiple *TIC236* gain-of-function mutations (TIC236GFs). Despite the low sequence identity between TIC236 and bacterial TamB, each mutated TIC236GF residue is conserved in TamB. Consistently, a *tic236*- knockdown mutant exhibited multiple lesion phenotypes similar to *crl*, indicating a shared functionality of CRL and TIC236. Ensuing reverse genetic analyses revealed genetic interaction between CRL and SP1, a RING-type ubiquitin E3 ligase, as well as with the plastid protease FTSH11, which function in TOC and TIC protein turnover, respectively. Loss of either SP1 or FTSH11 rescued *crl* mutant phenotypes to varying degrees due to increased translocon levels. Consistent with impaired plastid division exhibited by both *crl* and *tic236*-knockdown mutants, CRL interacts with the transit peptides of proteins essential in plastid division, and TIC236GF mutant proteins reinforce their import via increased TIC236 stability. Overall, our data shed new light on the links between plastid division, plant stress response and plastid protein import. We have also isolated and characterized the first GF mutants exhibiting increased protein import efficiency, which may inspire chloroplast engineering for agricultural advancement.

---

Chloroplasts evolved from a gram-negative cyanobacterial endosymbiont, with most cyanobacterial genes having been transferred to the host plant genome. Therefore, thousands of nuclear-encoded chloroplast proteins are post-translationally imported into chloroplasts, orchestrated by outer and inner envelope membrane (OEM and IEM) translocons, respectively termed TOC and TIC. Although an array of translocon proteins has been identified^1,2^, it had remained unclear how TOC and TIC accurately coordinate protein import across the two envelope membranes separated by an intermembrane space. Chen et al. (2018) shed some light on this question by discovering the TIC236 component, a homolog of the bacterial TRANSLOCON ASSEMBLY MODULE B (TamB)^3^. TIC236 is an integral IEM protein associated with TIC components. Its C-terminal domain, located in the intermembrane space, directly interacts with the N-terminal polypeptide transport-associated (POTRA) domains of TOC75-III (hereafter TOC75), the channel protein in the TOC complex. The fact that TOC75 contains three POTRA domains like TamA^4–6^ and that TIC236 contains a TamB-like domain (annotated as DOMAIN OF UNKNOWN FUNCTION 490) at the C terminus^3,7^ clearly reflects their bacterial origins^5^. Like *toc75* mutants, *tic236* null mutants display embryonic lethality, indicating that the function of TIC236 is indispensable in plants. Consistently, chloroplasts isolated from viable *tic236*-knockdown (*kd*) mutants exhibit significantly deficient protein import capability^3^. These findings strongly suggest that the ‘bacterial exit route evolved into an entry path in plants’^8^.

Plastid division occurs in developing cells to ensure an optimal number of plastids is in place before cell division, requiring the import of a suite of plastid-division machinery (PDM) proteins. The loss of any vital PDM elements results in gigantic plastids and a drastically reduced plastid number per cell^9^. Unexpectedly, several Arabidopsis mutants deficient in plastid division, including *crumpled leaf* (*crl*), develop foliar cell death^10^, resembling lesion-mimicking mutants (LMM) that exhibit a light-dependent hypersensitive response-like cell death^11,12^. Like LMM, *crl* and other plastid division mutants constantly upregulate immune-related genes^10,13^. The gigantic chloroplasts of *crl* mutants also induce an abnormal cell cycle, with increased endoreduplication activity leading to stunted growth^14^. Previous studies have indicated that autoimmune responses, abnormal cell cycle, and growth inhibition are likely mediated by a process called retrograde signaling, i.e., signaling from the gigantic chloroplasts back to the nucleus^10,13,14^.

CRL is a nuclear-encoded chloroplast OEM protein. Its short N-terminal region resides in the intermembrane space, followed by a transmembrane domain and then a chromophore lyase CpcT/CpeT domain characterized from a cyanobacterial CpcT bilin lyase^15^. Although the lyase domain retains phycocyanobilin-binding aptitude^16^, there is no apparent correlation between phycocyanobilin-binding ability and *crl*-induced lesions in Arabidopsis^17^, indicating that CRL has gained a divergent function.

## Dominant *TIC236* gain-of-function mutations abolish *crl*-induced lesions

To explore the function of CRL, we performed an ethyl methanesulfonate (EMS) mutagenesis screen to find suppressors of *crl* (*spcrl*). The mutagenized Arabidopsis M_2_ seeds were germinated on soil, and plants showing a wild-type (WT)-like phenotype were selected for further analyses. Among ∼24,000 M_2_ plants, we found two robust *spcrl* mutants, namely *spcrl1* and *spcrl2*, whose visible phenotype is nearly indistinguishable from that of WT plants, as well as one (*spcrl3*) that displayed weaker suppressing efficacy, especially in terms of restoring plastid division (Fig. 1a and Extended Data Fig. 1). Whole-genome sequencing of genomic DNA isolated from each of these *spcrl* mutants identified putative causal missense mutations in *TIC236*, specifically TIC236^D1212N^ in *spcrl1*, TIC236^G1250E^ in *spcrl2*, and TIC236^G1489R^ in *spcrl3* (Fig. 1b). Despite the low protein sequence identity (∼29%) between Arabidopsis TIC236 and *Escherichia coli* TamB, all three mutated residues are conserved in TamB (Extended Data Fig. 2) and across various plant species (Fig. 1c), highlighting their importance. The phenotypes of genetically isolated *tic236*(*D1212N*), *tic236*(*G1250E*), and *tic236*(*G1489R*) single mutants proved similar to WT plants (Fig. 1d). To distinguish these gain-of-function (GF) mutants from *tic236*-*kd* mutants, i.e., *tic236-2* (SAIL104-F07, *Columbia* ecotype) and *tic236-3* (RIKEN PST00216, *Nossen* ecotype)^3^, we re-named *tic236*(*D1212N*) as *tic236-4gf, tic236*(*G1250E*) as *tic236-5gf*, and *tic236*(*D1489R*) as *tic236-6gf*, respectively (Fig. 1d). Next, we crossed the isolated single mutant plants with the *crl* mutant to generate all possible genotypes in F2 siblings. PCR-based genotyping of the GF mutants confirmed a dominant effect of the *TIC236-4GF* and *TIC236-5GF* mutations and a less dominant effect of *TIC236-6GF* in rescuing the *crl* phenotypes (Fig. 1e and f). Since *crl* causes constitutive expression of stress-related genes^10,13,14^, next we conducted global gene expression profiling of *spcrl1* relative to the *crl* mutant. Our results confirmed that the *TIC236-4GF* mutation is epistatic to *crl* (Fig. 1g, Extended Data Fig. 3, and Supplementary Table 1).

**Fig. 1.**
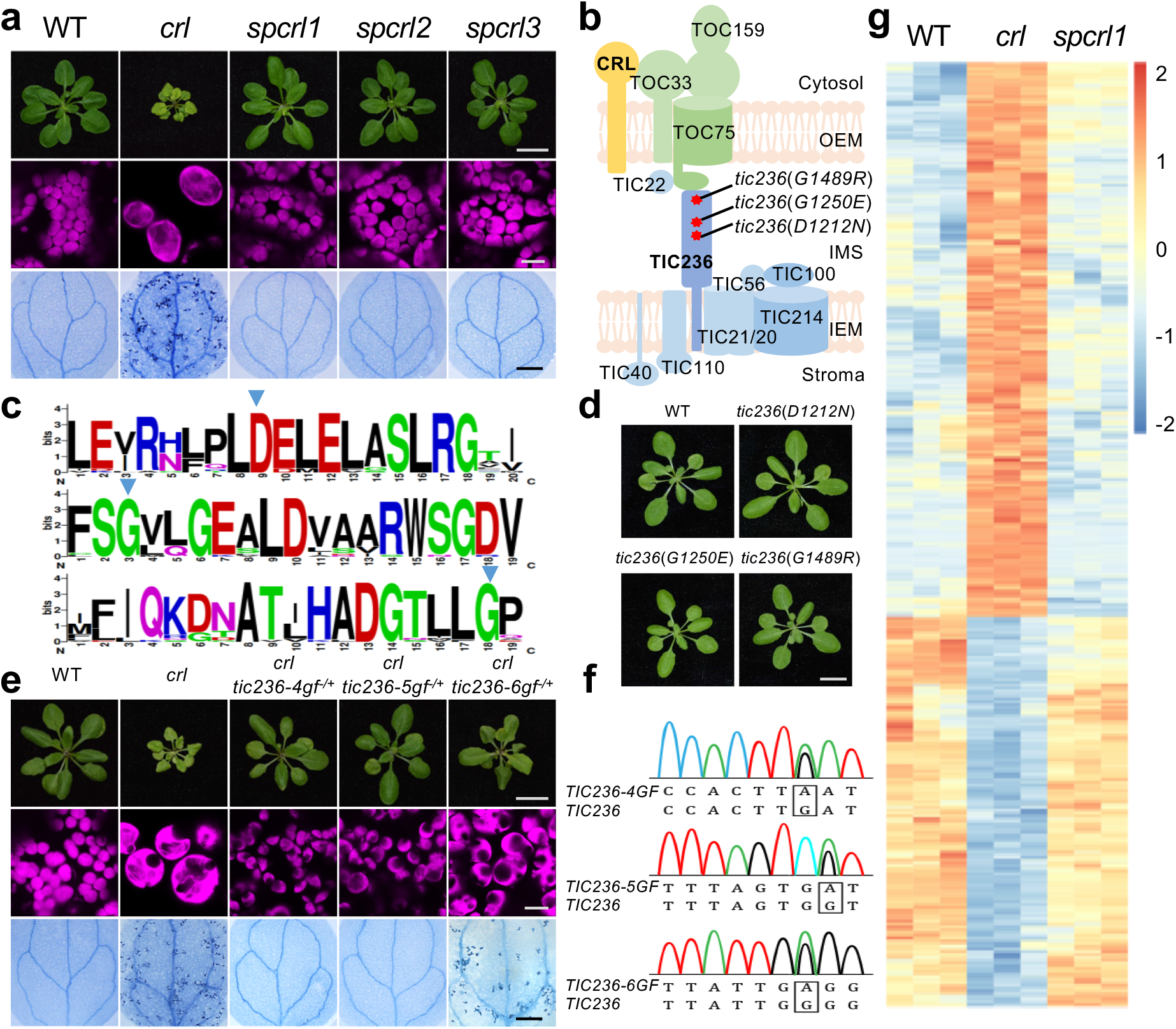
Identification of *TIC236* gain-of-function mutations in *spcrl* mutants. **a**, The micro and macroscopic phenotypes of the *crl* suppressors *spcrl1*, *spcrl2*, and *spcrl3*. Top: Representative images of 21-d-old plants grown on soil (scale bar, 1 cm). Middle: Confocal images representing chlorophyll autofluorescence (magenta) of mesophyll cells. Bottom: Cell death in 10-d-old cotyledons, as visualized by trypan blue (TB) staining (scale bar, 0.5 mm). **b**, Whole-genome sequencing reveals the causative missense mutations in a single genetic locus encoding the TIC236 protein, as indicated by red asterisks. **c**, The TIC236-like protein sequences from 27 plant species reported in Phytozome v12.1 were aligned using ClustalW. The three mutated residues in *spcrl* mutants are indicated with blue arrowheads. Aligned regions were visualized using WEBLOGO. **d**, Plant phenotypes of genetically isolated single mutants are shown (scale bar, 1 cm). **e**, Different levels of dominance for the D1212N (TIC236-4GF), G1250E (TIC236-5GF), and G1489R (TIC236-6GF) mutations are shown in each *tic236-4gf*, *tic236-5gf*, and *tic236-6gf* heterozygote (-/+) in the *crl* null mutant background (scale bars, 1 cm, 10 μm, and 0.5 mm, respectively). **f**, Sanger DNA sequencing result of *TIC236* mutations in the plants shown in (**e**). The heterozygous *TIC236* mutations are indicated with black boxes. **g**, Heatmap showing the differentially expressed genes in *crl* versus WT and *spcrl1* plants. The genes with at least a two-fold change in expression and a false discovery rate of less than 0.05 were selected. The colors of the heatmap represent the z-scores ranging from –2.0 (blue) to 2.0 (red).

### CRL is associated with OEM translocon components

Our discovery of the *TIC236GF* mutations among the *crl* suppressors led us to consider CRL as a probable translocon-associated factor. Accordingly, we co-immunoprecipitated biologically active GFP-tagged CRL and its accompanying proteins using GFP-conjugated magnetic Dynabeads (Extended Data Fig. 4a and b). Eluted proteins were trypsin-digested and subjected to mass spectrometry (MS) analysis, revealing a total of 187 proteins (detected at least twice in *CRL-GFP crl* but not in *GFP* plants) (Supplementary Table 2), including TOC75, TOC34, TOC132 and TIC110, but not TIC236 (Extended Data Fig. 4c). Another eight import-related proteins—including TOC33, TOC159, TOC120, TIC20, and TIC214—were identified as putative albeit less significant CRL-associated proteins (Supplementary Table 3), which were only detected once in *CRL-GFP crl* and not in *GFP* samples. To date, we have not detected endogenous CRL or TIC236 in WT plants via our MS-based label-free chloroplast proteome assay, indicating their lower abundances and/or instability. Nonetheless, we validated the CRL-TOC interaction by means of coimmunoprecipitation-Western blot analysis (Fig. 2a). We then chose TOC33 and TOC34 to verify a direct interaction with CRL *in vivo* using a bimolecular fluorescence complementation (BiFC) assay. Whereas no Venus fluorescence signal was detected when the N-terminal half of Venus fluorescence protein (nV) alone and C-terminal Venus (cV)-tagged TOC33 or TOC34 were co-expressed in *Nicotiana benthamiana* leaves, co-expression of either cV-TOC33 or cV-TOC34 with CRL-nV generated apparent Venus fluorescence signal in chloroplast envelopes (Fig. 2b). Coimmunoprecipitation and subsequent immunoblot analyses further confirmed these interactions (Fig. 2c).

**Fig. 2.**
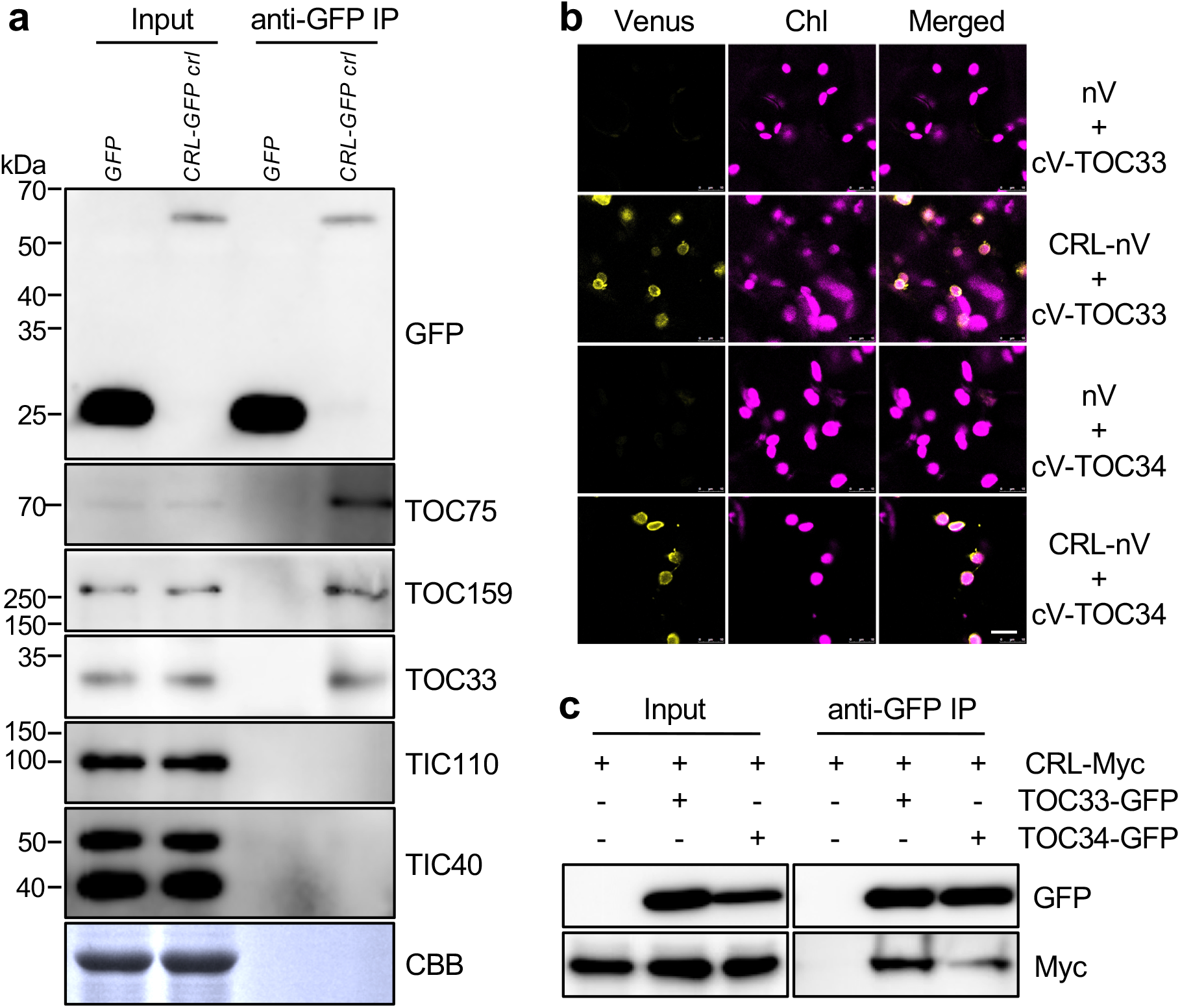
CRL is associated with TOC components. **a**, Co-IP/immunoblot results of CRL-TOC interactions. Arabidopsis stable *GFP* and *CRL-GFP crl* transgenic plants were used. Total proteins were extracted from 5-d-old seedlings and then free GFP and CRL-GFP were pulled down. Subsequent immunoblot analysis was conducted using the indicated TOC and TIC antibodies. Coomassie blue (CBB) staining of the SDS-PAGE gels is shown as a loading control (for input samples). **b**, BiFC analysis. N-terminal Venus (nV)-fused CRL and C-terminal Venus (cV)-fused TOC33 (or TOC34) were transiently coexpressed in *N. benthamiana* leaves. The empty nV vector was used as a negative control. The representative Venus signals, chlorophyll autofluorescence (Chl) signals, and their merged images are shown. Scale bar, 10 μ **c**, Co-IP/immunoblot analysis. Different construct combinations, such as CRL-Myc alone, CRL-Myc and TOC33-GFP, or CRL-Myc and TOC34-GFP, were transiently overexpressed in *N. benthamiana* leaves. After 2 days, GFP-Trap magnetic beads were used to enrich the target and its associated proteins. Subsequent immunoblot analyses were carried out by using anti-GFP and anti-Myc antibodies.

### Reduced translocon turnover substantially rescues the *crl* phenotypes

TOC components undergo proteolysis via the ubiquitin-proteasome system (UPS) in a process referred to as chloroplast-associated protein degradation (CHLORAD)^18,19^. Together, the OEM-spanning RING-type ubiquitin E3 ligase, i.e., suppressor of *ppi1* locus 1 (SP1), the TOC75-like protein SP2 that lacks POTRA domains, and the cytosolic AAA+ chaperone CDC48 direct CHLORAD^20^ (Fig. 3a). Loss of SP1 increases the amount of TOC components, whereas SP1 overexpression (oxSP1) reduces them. SP1-mediated TOC degradation confers on plants tolerance to oxidative stress by decreasing the levels of reactive oxygen species (ROS, byproducts of photosynthesis), which arises from decreased import of photosynthesis-associated proteins and thus demonstrating the physiological importance of SP1^21^. If CRL protein functions in protein import, *sp1*-driven TOC accumulation may attenuate the *crl*-induced phenotypes. Indeed, we found that loss of SP1 largely rescued the *crl* phenotypes (Fig. 3b, d, and Extended Data Fig. 5), just as it considerably rescued the *ppi1* mutant lacking TOC33^20^. This finding prompted us to also generate *crl ftsh11* double-knockout mutants. FTSH11 is a plastid metalloprotease implicated in TIC40 turnover^22^. FTSH11 also physically interacts with CHAPERONIN 60 (CPN60), the activity of which is required for precursor protein maturation after excision of the transit peptide and plastid division^22,23^. Similar to SP1 that is required under oxidative stress conditions, FTSH11 plays a vital role in thermotolerance^24,25^. Loss of FTSH11 significantly rescued the *crl* phenotypes, including those of plastid ultrastructure and division, as well as cell death (Fig. 3c, d, and Extended Data Fig. 6a and b). Moreover, loss of either SP1 or FTSH11 increased the levels of translocon components in both the *crl* mutant and WT (Fig. 3e and f). This reverse genetic approach using *sp1* and *ftsh11* mutants further reinforces the evidence for CRL being a translocon-allied component.

**Fig. 3.**
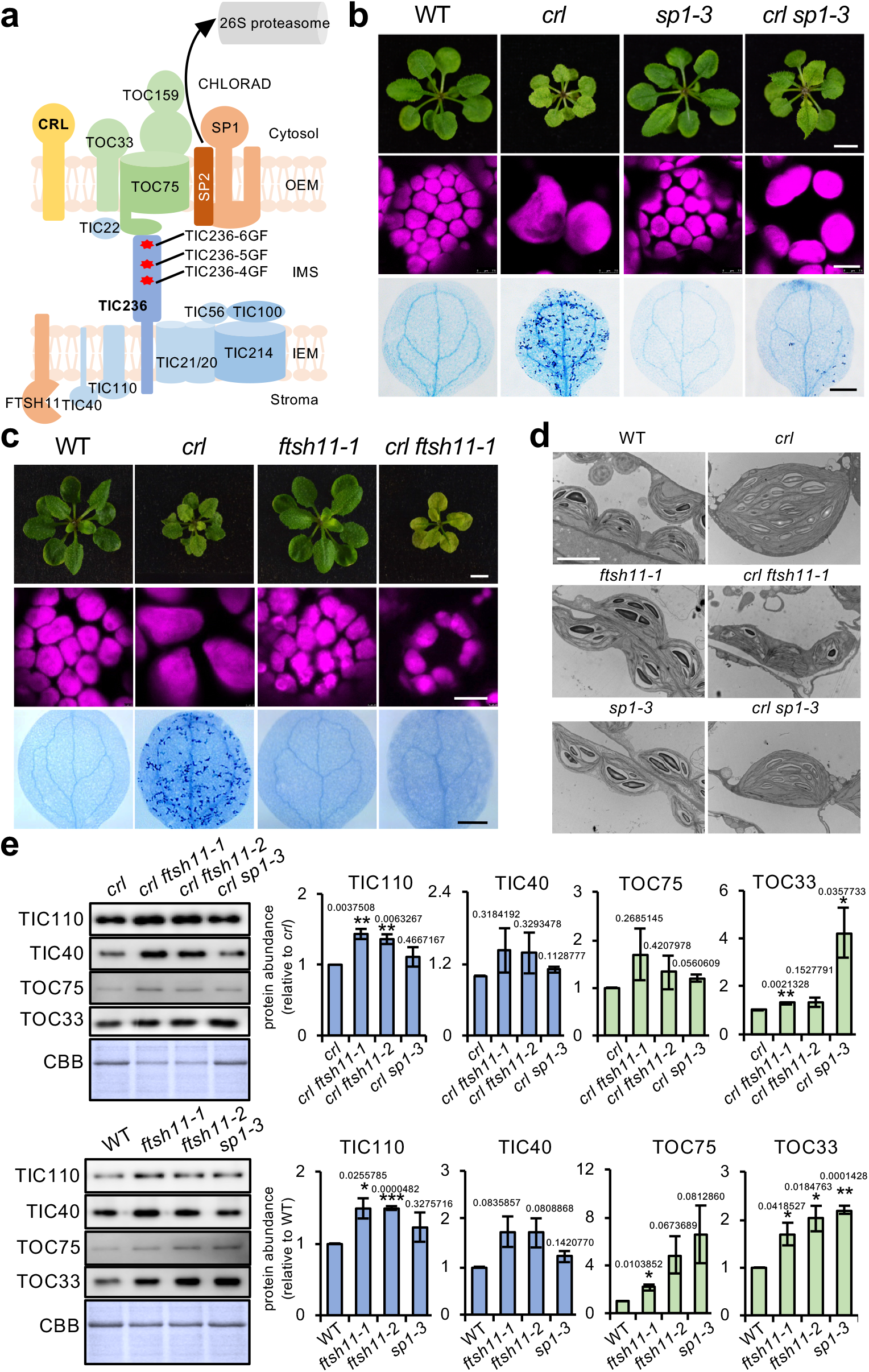
Loss of either SP1 or FTSH11 significantly rescues the *crl* phenotypes. **a**, SP1- and FTSH11-mediated TOC and TIC turnover^18,20,22^. **b**-**c**, Images representing 21-d-old plants (top), chloroplasts in mesophyll cells (middle), and cell death in cotyledons (bottom) are shown (Scale bars: top, 5 mm; middle, 10 µm; bottom, 0.5 mm). **d**, The chloroplast ultrastructure of 5-d-old seedlings was monitored by transmission electron microscopy (scale bar, 5 µm). **e**, From the same plant materials as in (**b**) and (**c**), equal amounts of total protein were separated on SDS-PAGE gels and immunoblotted with the antibodies as indicated to detect the relative abundance of TIC and TOC proteins. The quantified protein abundance is shown as mean ± SD of three independent biological repeats after normalization to *crl* or WT of the same experiments. All *P*-values are from Student’s *t*-tests (two-tailed). \**P* < 0.05, \*\**P* < 0.01, \*\*\**P* < 0.001. CBB staining of the SDS-PAGE gels is shown as a loading control.

### A *TIC236* knockdown mutant also exhibits cell lesion and defective plastid division phenotypes

Next, we compared the phenotypes of *crl* and the *TIC236* knockdown mutant *tic236-2*. Remarkably, all *crl* phenotypes, such as growth retardation, localized cell death, and the plastid division defect, were recapitulated in *tic236-2* plants (Extended Data Fig. 7a). The *tic236-2* mutant also exhibited aplastidic guard cells (only one chloroplast per guard cell pair), which resulted from impaired plastid division during stomatal biogenesis^26^. It is important to note that whereas *CRL* mutation causes persistently impaired plastid division, *tic236-2* mutant plants exhibit significantly uneven numbers and sizes of plastids per cell (Extended Data Fig. 7b and c), perhaps because different cellular protein levels of TIC236 heterogeneously impede plastid division. For instance, if cells contain TIC236 above a certain threshold level, leading to import of sufficient amounts of PDM preproteins, then plastid division would be indistinguishable from that displayed by WT cells. However, if levels of TIC236 are below a certain threshold, then plastid division might be significantly impaired. Interestingly, *tic236-2* exhibited more cell death relative to that observed in the *crl* mutant, indicating that a combination of reduced general import capability and the plastid division defect may exacerbate cell death. The *crl*-induced defect in plastid division was epistatic to *tic236-2*, as only gigantic chloroplasts were observed in *crl tic236-2* double mutant plants (Extended Fig. 7a). This epistatic relationship reveals a fundamental role for CRL in plastid division and a synergistic impact of *TIC236* knockdown (e.g., via general import) on *crl*-induced growth retardation and cell death (Extended Fig. 7a).

### GF mutations stabilize TIC236 and enhance protein import aptitude

Our above described results suggest that *TIC236GF* mutations suppress *crl*-induced lesions via an enhanced protein import rate. Accordingly, we further examined this possibility by means of an *in vitro* import assay. We excluded the *crl* mutant itself because of its huge chloroplast size that impedes isolation of intact chloroplasts. [^35^S]methionine-labeled preproteins, including CASEIN LYTIC PROTEINASE C1 (prCLPC1, also known as prHSP93), TRANSLOCON AT THE INNER ENVELOPE MEMBRANE OF CHLOROPLASTS 40 (prTIC40), OXYGEN EVOLVING COMPLEX SUBUNIT 23 kD (prOE23), and the PDM components FILAMENTING TEMPERATURE-SENSITIVE Z (FTSZ) homolog (prFTSZ) 2-1 and prFTSZ2-2 were incubated with chloroplasts isolated from each genotype. The preproteins prOE23, prHSP93, and prTIC40 represent proteins that reside in thylakoid, stroma, and IEM, respectively. Consistent with a previous report^3^, *tic236-2* chloroplasts displayed significantly impaired import of all tested preproteins (Fig. 4a and b). In contrast, we observed enhanced preprotein import for *tic236-4gf* and *tic236-5gf* chloroplasts, and less so for *tic236-6gf* chloroplasts, in agreement with the respective plant phenotypes (Fig 1a, e and Extended Data Fig. 1). It is possible that the increase in *TIC236-6GF*-caused import was not adequately detected via *in vitro* import assay, which assesses import rate within an extremely short time period. However, the slight increase in protein import induced by *TIC236-6GF* throughout plant development may be sufficient to partially surpress the multiple defects caused by loss of CRL.

**Fig. 4.**
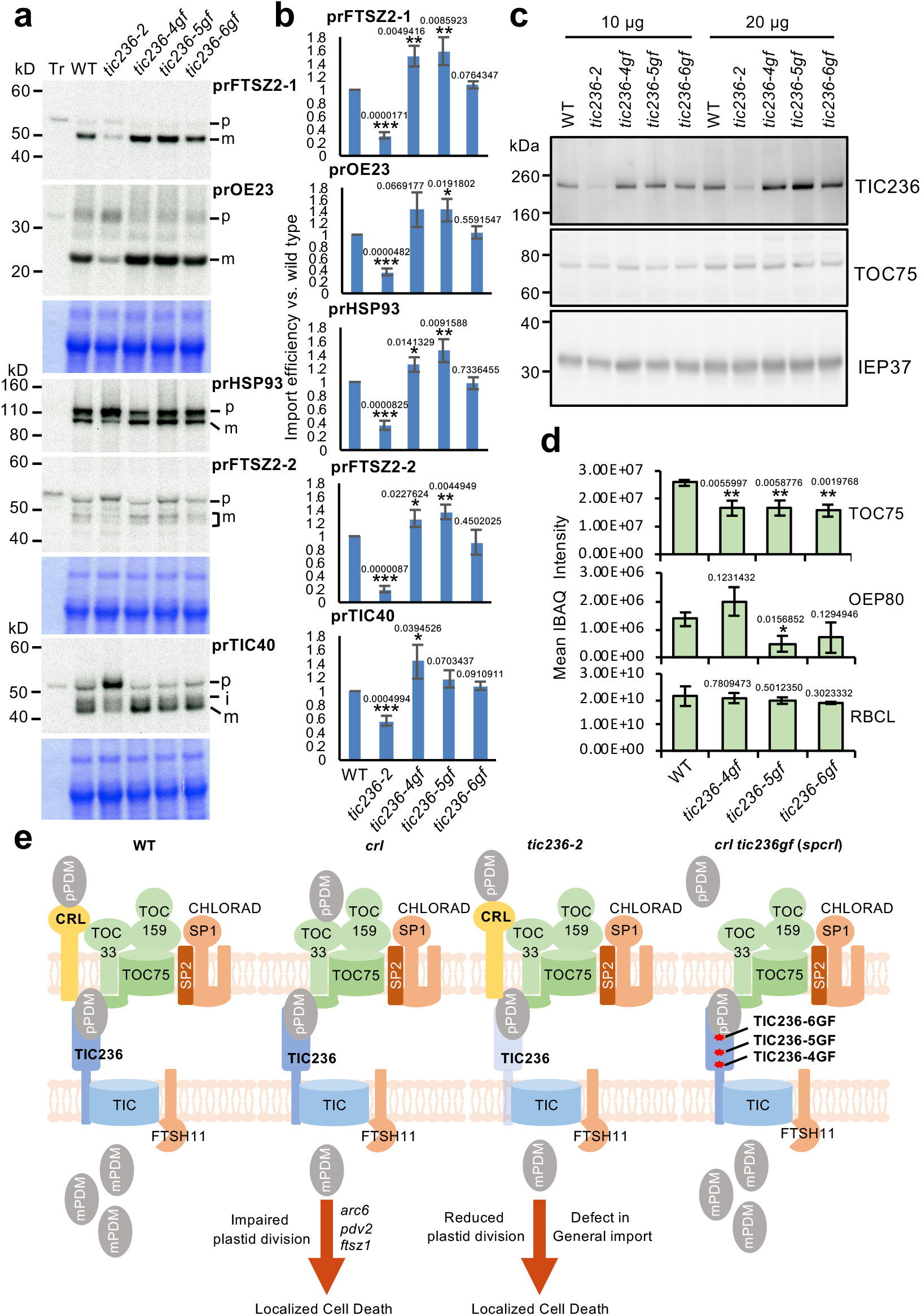
*TIC236GF* mutations rescue the *crl* mutant by stabilizing TIC236 proteins. **a**, [^35^S]Met-labeled preproteins (Tr) were imported into chloroplasts isolated from 14-d-old plants of WT and the four *tic236* mutant alleles in 3 mM ATP at room temperature for 10 min. prFtsZ2-1 and prOE23 were co-imported, as was prHsp93 and prFtsZ2-2. Re-isolated intact chloroplasts were analyzed by SDS-PAGE and the gels were stained with CBB and dried for fluorography. Equal amounts of proteins were loaded in each lane of the same gel, except for the Tr lane. The region around the chlorophyll *a/b* binding protein in the CBB-stained gels is shown below the fluorograph as a loading control. **b**, Imported mature proteins were quantified and normalized to that of the WT from the same gel and further corrected by the amount of the chlorophyll *a/b* binding protein in each lane. Data shown are mean ± SD (n=3). *P*-values were obtained from two-tailed Student’s *t*-tests. \**P* < 0.05, \*\**P* < 0.01, \*\*\**P* < 0.001. **c**, Chloroplasts isolated from the indicated genotypes were analysed by SDS-PAGE, followed by blotting onto nitrocellulose blotting membranes (Amersham Protran), and then hybridized with antibodies against proteins indicated at right. The dilutions used were: anti-Tic236 1:1000; anti-Toc75 1:6000; and anti-IEP37 1:4000. **d**, Label-free protein quantitation. iBAQ intensities are shown as mean ± SD (n=3). Ribulose-bisphosphate carboxylase large-chain (RBCL) was chosen as the control. All *P*-values are from two-tailed Student’s *t*-tests. \**P* < 0.05, \*\**P* < 0.01. P < 0.05. **e**, In WT, prPDM are imported through the CRL-TIC236-translocon module, enabling plastid division. Either CRL loss or TIC236 knockdown compromises plastid division and induces cell death despite the core translocon complex remaining intact. Similar phenotypes were observed in the canonical plastid division mutants *arc6*, *pdv2*, and *ftsz1*^10,13^. The TIC236GF mutations overcome CRL deficiency by reinforcing prPDM import. Reduced SP1-driven CHLORAD activity or loss of FTSH11 protease also rescues the *crl* phenotypes.

Assuming that the CRL-TIC236 module is required for importing PDM, we also examined the relative import rate of the vital PDM components FTSZ2-1 and FTSZ2-2. We observed enhanced import rates of both those proteins into *tic236-4gf* and *tic236-5gf* chloroplasts, and again less so in *tic236-6gf* chloroplasts (Fig. 4a, b), further evidencing a function for the CRL-TIC236 module in importing PDM for plastid division. Accordingly, we hypothesized that *TIC236GF* mutations promote preprotein import by stabilizing the TIC236 protein, which may compensate for the lack of CRL protein in chloroplasts. Indeed, we found that *TIC236-4GF* and *TIC236-5GF* mutations increased the steady-state levels of cognate mutant proteins compared to TIC236 in WT plants (Fig. 4c).

Impaired plastid division has not been reported for the few translocon mutants documented as viable, except for mutants of the chaperonin CPN60 complex that is required for protein import and thylakoid membrane protein insertion^23,27,28^. Since PDM import is a prerequisite for plastid division, we presumed that the CRL-TIC236 module might import some PDM components, whereas the well established TIC236- harboring TOC/TIC complex acts in universal import, as evidenced by the embryonic lethality of the *tic236* knockout mutant. If this notion is correct, CRL might interact with the transit peptides of PDM components. BiFC assays confirmed interactions at the OEM of CRL with the transit peptides of FTSZ2-1 and FTSZ2-2 but not with those of RBCS and FTSZ1 (Extended Data Fig. 8a), suggesting its specificity towards certain PDM proteins. However, coexpression of CRL with the mature form (lacking the transit peptide) of either FTSZ2-1 and FTSZ2-2 resulted in abnormal Venus signals (foci and rod shapes, Extended Data Fig. 8b).

### TIC236 is functionally divergent from its ancestral TamB protein

TIC236 homologs have been identified from all plant species in which they were sought. In rice, loss of SUBSTANDARD STARCH GRAIN 4 (SSG4), an ortholog of maize DEFECTIVE KERNEL 5 (DEK5) and Arabidopsis TIC236, causes plastid abnormality^29^. In maize, DEK5 inactivation impairs both plastid division and plastid envelope proteostasis, e.g., reduced or absent protein levels of translocon components and inorganic phosphate transporters^30^, so DEK5 appears to have retained its TamB functionality, mostly related to biogenesis of outer membrane proteins. In particular, levels of the OMP85-type β-barrel proteins TOC75 and OEP80 were diminished in *dek5* mutants^30^. If TIC236 functions in OEM protein biogenesis, we might expect to see reduced levels of translocons in *tic236-2* and increased levels in *tic236-gf* mutants, respectively. However, the *tic236-2* mutant does not exhibit reduced levels of TOC75^3^. Furthermore, all three of our *tic236-gf* mutant lines displayed comparable levels of TOC75 protein relative to WT plants (Fig. 4c). To further verify that result, we conducted label-free quantitative proteomics analyses using isolated chloroplasts of WT and *tic236-gf* mutants and found that levels of most of the TOC/TIC components in the *tic236-gf* mutants remained unchanged relative to WT (Fig. 4d and Supplementary Table 4).

CRL is believed to function in a ROS-triggered chloroplast-to-nucleus retrograde signaling pathway or as a putative PDM component in plastid division^10,14,15^. However, our study using combined biochemical and forward/reverse genetic approaches has revealed its mutual functionality with the TIC236 protein. Since the *tic236* null mutant displays embryonic lethality^3^, whereas the *crl* mutant is viable despite multiple lesions (Fig. 1a), unlike CRL, the function of TIC236 in plastid translocons must be indispensable. The gigantic plastids in both the *crl* and *tic236-2* mutants (Extended Data Fig. 7), as well as the verified CRL-PDM interaction (Extended Data Fig. 8a), imply that the CRL-TIC236 module imports PDM proteins at the early phase of cell development. The observed cell death and chloroplast division phenotypes displayed by the *crl* and *tic236-2* mutants should be investigated in yet other translocon mutants to gain further insights into the passenger specificity of the CRL-TIC236 module, as well as into possible spatio-temporal heterogeneity of translocon complexes and its biological relevance. Importantly, our findings also open up a new research avenue linking chloroplast dysfunction (especially certain import pathways) to activation of autoimmune responses (Fig. 4e). In summary, we have reported herein that: (i) CRL is a translocon-associated component, absence of which impairs plastid division; (ii) knockdown of *TIC236* induces multiple lesions and plastid division defects, as also observed in the *crl* mutant; (iii) three approaches for increasing translocon component abundance—knockout of either *SP1* or *FTSH11*, and gain-of-function *TIC236* mutations—all rescued *crl* plastid divison defects; and (iv) *TIC236GF* mutations may provide a strategy for engineering the translocon to enhance protein import efficacy.

## Materials and Methods

### Plant material and growth conditions

All Arabidopsis seeds used in this study are from the *Columbia-0* (*Col-0*) background. All seed stocks, including *ftsh11-1* (SALK_033047), *ftsh11-2* (SALK_012285), *sp1-3* (SALK_002099), and *tic236-kd* (*tic236-2*, SAIL_104-F07), were obtained from the Nottingham Arabidopsis Stock Centre (NASC). The *crl* null mutant (GABI_714_E08) and the transgenic *p35S::CRL-GFP crl* lines have been described previously^13,17^. Their genotypes were confirmed by PCR-based analyses using the corresponding primers (Supplementary Table 5). Seeds were sterilized in a 5% hypochlorite solution for 3 min, washed five times with sterilized water, and then sown on half-strength Murashige and Skoog (MS) medium (Duchefa Biochemie) with 0.5% (w/v) sucrose and 0.7% (w/v) agar. After stratification at 4 °C for 2 days (d) in the dark, the seeds germinated and grew under either continuous light (CL, 100 µmol m^-2^ s^-1^ at 22 ± 2 °C) or long-day (LD, 22 ± 2 °C, 100 µmol m^-2^ s^-1^ with a 16 h-light/8 h-dark photoperiod) conditions. *Nicotiana benthamiana* plants were grown under controlled LD conditions. Four-week-old *N. benthamiana* plants were used for all transient expression assays.

### EMS mutagenesis and whole-genome sequencing analysis

Under normal growth conditions, we screened the M_2_ progeny of ∼12,000 M_1_ *crl* seeds that had been initially treated with 0.4% (w/v) ethylmethanesulfonate (EMS, Sigma-Aldrich) for 8 h as described previously^31^. The *spcrl1*, *spcrl2*, and *spcrl3* mutants were chosen to generate mapping populations. To do that, each *spcrl* homozygote mutant plant was backcrossed to the parental *crl* mutant. From the F_2_ population, at least 50 plants exhibiting WT-like phenotypes were selected for genomic DNA extraction. Genomic DNA isolated from *crl* mutant plants was used as a control. Genomic DNA (1 μg) isolated using a DNeasy plant mini kit (Qiagen) was used for library construction. Sequencing was performed using a HiSeq2500 (Illumina) sequencer to generate 125 base pair (bp) paired-end reads as described in Li et al. (2020). The sequencing data were processed in SolexaQA^32^ and Cutadapt (v.1.3) software to remove low-quality regions and adapter sequences, respectively. Clean reads were mapped to the TAIR10 genome in BWA-MEM^33^ with default parameters. SNPs were called using the “mpileup” function of SAMtools^34^. Poor quality SNPs with a mapping quality <60 or with a depth <3 or >200 were filtered out using vcftools^35^. Candidate causal mutations were identified using the SHOREmap method^36^. The allele frequency and the regions containing a possible causal mutation were analysed using SHOREmap v3.0^37^. Mutations within the open reading frame (ORF) of target genes were considered as potential causative mutations. The mutations were further confirmed by sequencing PCR products using the respective primers listed in Supplementary Table 5.

### Coimmunoprecipitation and MS analyses

The cDNA (lacking stop codons) of *CRL*, *TOC33*, and *TOC34* were PCR-amplified from WT cDNA and subsequently cloned into pDONR221-Zeo entry vectors by means of a Gateway BP clonase reaction (Invitrogen). To generate C-terminal tag-fused constructs, the purified cDNAs were cloned into destination vectors, including pGWB605 for sGFP or pGWB617 for 4xMyc, by Gateway LR clonase reactions (Invitrogen). Each vector was transformed into *Agrobacterium tumefaciens* strain GV3101. For transient expression (or co-expression), the suspensions of agrobacteria carrying different constructs were infiltrated into healthy leaves of 28-d-old *N. benthamiana* plants and protein-protein interactions were analyzed after 48 h. Total protein was extracted using a protein extraction (PE) buffer [Tris-HCl 50 mM (pH 7.5), NaCl 150 mM, Glycerol 10% (v/v), DTT 10 mM, cOmplete protease inhibitor cocktail 1 tablet/50 ml (Roche), 1.1% (v/v) NP-40, EDTA 1 mM, Na_2_MoO_4_ 1 mM, Sodium fluoride (NaF) 1 mM, and Sodium orthvanidate (Na_3_VO_4_) 1.5 mM]. After diluting total protein samples to 1-2 l, μ 20 mg of total protein was incubated with 15 μl GFP-Trap^MA^ beads (Chromotek) at 4 °C for 2 h, and then washed four times with PE buffer. Finally, the remaining proteins on the washed beads were eluted with 2x SDS sample buffer at 95 °C for 10 min. The eluted proteins were subjected to 10% SDS-PAGE gels for immunoblot analyses using anti-GFP (Roche) and anti-Myc (Roche) antibodies.

For coimmunoprecipitation (Co-IP) and subsequent immunoblot analysis or Co-IP coupled to MS analysis, 14-d-old Arabidopsis rosette leaves were used. Rosettes were ground to a fine powder in liquid nitrogen and resuspended in IP buffer [HEPES 20 mM (pH 7.5), EDTA 2 mM, EGTA 2 mM, NaCl 100 mM, Glycerol 10% (v/v), Triton-X-100 0.2% (v/v), Na_3_VO_4_ 1 mM, NaF 20 mM, and cOmplete protease inhibitor cocktail 1 tablet/50 ml (Roche)] at 4 °C for 1 h. After quantification using a PierceTM BCA protein assay kit (Thermo Scientific), 10 mg of total protein was incubated with 50 μl Dynabeads Protein G (Thermo Scientific) that had been pre-conjugated with a monoclonal mouse anti-GFP antibody (Roche) at room temperature for 2 h. After washing five times, 10% of the beads were eluted with SDS sample buffer (2X) at 70 °C for 20 min, and then loaded on 10% SDS-PAGE gels for subsequent immunoblotting analysis.

In-gel digestion and ensuing MS analysis were performed according to a previous report^38^. The obtained mass spectra were compared against the TAIR10 non-redundant database using Mascot Server (v2.5.1). The parameter settings were set for Peptide mass tolerance at 20 ppm, fragment mass tolerance at 0.02 Da, and a maximum of two missed cleavages was allowed. The significance threshold for search results was set at P < 0.05, with an ion score cutoff of 15. Proteins detected at least twice in the CRL-GFP samples, but not in the GFP samples, were considered potential interacting proteins of CRL (Supplementary Table 2).

### BiFC assays

In addition to the full-length cDNA of *CRL*, *TOC33*, and *TOC34*, cDNA encoding the transit peptide (tp) or mature (m) regions (without the stop codon) of *RBCS*, *FTSZ1*, *FTSZ2-1*, and *FTSZ2-2* were cloned into pDONR221-Zeo entry vectors for BiFC analyses. Through Gateway LR reactions (Thermo Scientific), these cDNAs were cloned into the CaMV *35S* promoter (35S)*-*driven destination vector pDEST to generate tag-fused constructs. The full-length *CRL* cDNA (lacking the stop codon) was C-terminally fused with the N-terminal half of Venus (nV). The full-length *TOC33* and *TOC34* cDNAs were N-terminally fused with the C-terminal half of Venus (cV). cDNA corresponding to the transit peptides and mature regions of RBCS, FTSZ1, FTSZ2-1, and FTSZ2-2 were C-terminally fused with cV. As described previously^39^, the N-terminal 79 residues of RBCS were constructed as the transit peptide region. Transit peptide regions of FTSZ1 (1-90), FTSZ2-1 (1-48), and FTSZ2-2 (1-50) were predicted by the ChloroP1.1 server^40,41^. Then, tp-deleted and start codon-appended RBCS, FTSZ1, FTSZ2-1, and FTSZ2-2 constructs were generated. Destination plasmids CRL-nV, cV-TOC33, cV-TOC34, tpRBCS-cV, tpFTSZ1-cV, tpFTSZ2-1-cV, tpFTSZ2-2-cV, mRBCS-cV, mFTSZ1-cV, mFTSZ2-1-cV, and mFTSZ2-2-cV were transformed into *Agrobacterium* strain GV3101. The *Agrobacterium* cultures were diluted to an OD_600_ of 1.0, then resuspended and washed using infiltration solution [MES 10 mM (pH 5.6), MgCl_2_ 10 mM, and Acetosyringone 150 μM]. Mixtures of the selected strains were infiltrated into 28-d-old *N. benthamiana* leaves. After transient expression for 48 h, Venus fluorescence was analysed using confocal laser scanning microscopy (TCS SP8, Leica).

### Protein extraction and immunoblot analysis

Plant leaf tissues were ground to fine powder in liquid nitrogen and resuspended in IP buffer at 4 °C for 1 h. Protein concentration was determined using a Pierce^TM^ BCA protein assay kit (Thermo Scientific). Equal concentrations of protein samples were mixed with 4x SDS loading buffer, denatured at 95 °C for 5 min, and loaded into 10% SDS-PAGE gels. GFP-tagged proteins were detected with a mouse anti-GFP antibody (1:5000; Roche). Immunoblot results were quantified using Image J software (v1.8.0).

### *In vitro* protein import assay

[^35^S]Met-labeled preproteins were *in vitro*-transcribed/translated using the TNT-coupled wheat germ or reticulocyte lysate system and SP6 or T7 RNA polymerase (Promega). Growth of Arabidopsis seedlings (for 14 days on MS agar media with 2% sucrose), Arabidopsis chloroplast isolation, and import of [^35^S]-labeled preproteins into isolated chloroplasts were performed as described previously^42^. Accession numbers for preproteins are: Pea prTIC40 (AY157668), Pea prHSP93 (L09547), Arabidopsis prOE23 (At1g06680), Arabidopsis prFTSZ2-1 (At2g36250), and Arabidopsis prFTSZ2-2 (At3g52750).

### Chloroplast size analysis

Leaf petioles of 14-day-old seedlings grown on MS medium were excised and fixed in 3.5% glutaldehyde and then prepared for imaging with differential interference contrast microscopes as described previously^43^.

### Cell death determination

Cell death was assessed via Trypan blue (TB) staining as described previously^13^. The TB-stained plants were preserved in 10% (v/v) glycerol. Imaging was conducted using a TCS SP8 microscope (Leica Microsystems) and further processed using Leica LAS software (v4.2.0, Leica Microsystems).

### Microscopic analyses

Venus and chlorophyll autofluorescence signals were monitored under a confocal microscope at 520-600 nm of the emission spectrum with an excitation wavelength of 514 nm (Leica TCS SP8). Representative images were processed using Leica LAS AF Lite software (v2.6.3, Leica Microsystems). Cotyledons of 5-d- and 10-d-old seedlings (before and after cell death, respectively) were mostly used for imaging GFP and chlorophyll autofluorescence unless otherwise indicated.

For transmission electron microscopy, cotyledons of 5-d-old seedlings were detached, pre-fixed, and then rinsed three times using 0.1 M PBS buffer as described previously^13^. Then the samples were post-fixed overnight in 1% (v/v) osmic acid at 4 °C, washed three times with 0.1 M PBS buffer, dehydrated using a gradient ethanol-acetone series, before being embedded in Spurr’s resin. Ultrathin resin sections (70 nm) were cut using a diamond knife on a Leica UC7 ultramicrotome, mounted on copper grids, and stained with 2% (w/v) uranyl acetate and 0.5% (w/v) lead citrate. The stained sections were monitored and photographed using a H7700 transmission electron microscope (Hitachi).

### RNA-seq library construction and data analysis

RNA-seq analysis was carried out as described previously^44^. Total RNA was extracted from three independent biological replicates of 5-d-old Arabidopsis seedlings of the *crl*, *spcrl1*, and WT using the RNeasy Plant Mini Kit (Qiagen). The isolated RNA was subjected to on-column DNase digestion according to the manufacturer’s instructions. A Nano Photometer spectrophotometer (IMPLEN) was used to verify RNA purity. A Qubit RNA Assay Kit and a Qubit 2.0 Fluorometer (Life Technologies) were employed to determine RNA concentration. An RNA Nano 6000 Assay Kit and Bioanalyser 2100 system (Agilent Technologies) were used to assess RNA integrity for RNA-seq analyses. RNA-seq libraries were built using the NEBNext Ultra Directional RNA Library Prep Kit for Illumina (New England Biolab) based on the manufacturer’s instructions. The RNA-seq libraries were sequenced on an Illumina HiSeq 2500 platform to generate 100 bp paired-end reads. The raw sequencing data were processed in SolexaQA (v2.2) to extract pair reads and to remove low-quality reads. The clean reads were mapped to the Arabidopsis genome (TAIR10) using TopHat^45^. After mapping, the Python-based software HTseq-count was used to extract raw counts of annotated genes. Differentially expressed genes (DEGs) were identified using the R package edgeR, which uses counts per gene in different samples and performs data normalization using the trimmed mean of M-values (TMM) method^46^. The gene expression data were normalized to transcripts per million (TPM) according to the total number of mapped clean reads in each library. Genes with at least a two-fold change in expression and a false discovery rate of less than 0.05 were deemed to be differentially expressed.

### RNA extraction and quantitative RT-PCR

Total RNA was prepared using the FastPure Plant Total RNA Isolation Kit (Vazyme). RNA (1 µg) was treated with RQ1 RNase-Free DNase (Promega) and reverse-transcribed using HiScript II Q RT SuperMix for qPCR (+gDNA wiper) (Vazyme) according to the manufacturer’s instructions. Quantitative RT-PCR was performed using ChamQ Universal SYBR qPCR Master Mix (Vazyme) and a QuantStudio^TM^ 6 Flex Real-Time PCR System (Applied Biosystems). Transcript abundances were calculated using the delta-Ct method^47^ and normalized to the transcript levels of *ACTIN2* (*AT3G18780*). The primers used in this study are listed in Supplemental Table 5.

### Chloroplast isolation and MS analysis

Chloroplasts were isolated from 21-d-old plants as described previously^48,49^. Rosette leaves were homogenized in chloroplast isolation buffer [50 mM HEPES-KOH (pH 8.0), 5 mM MgCl_2_, 5 mM EDTA (pH 8.0), 5 mM EGTA (pH 8.0), 10 mM NaHCO_3_, and 0.33 M D-sorbitol supplemented with one tablet (per 50 mL) of cOmplete protease inhibitor cocktail (Roche)] using a Waring blender. After filtering through four-layer Miracloth, the homogenate was centrifuged at 400 × g for 8 min at 4 °C. The pellets were suspended using chloroplast isolation buffer and added onto a two-step Percoll gradient (40:80%). After centrifugation, the enriched chloroplasts between the two Percoll steps were carefully collected and washed twice using HS buffer [50 mM Hepes-KOH (pH 8.0) and 0.33 M D-sorbitol]. The chloroplasts were resuspended in guanidine hydrochloride buffer [6 M guanidine hydrochloride and 100 mM Tris (pH 8.5)]. The resuspension was sonicated in an ice bath for 1 min with a pulse of 3 sec on and 5 sec off, followed by heating at 95 °C for 5 min, and centrifugation at 15000 rpm for 30 min at 4 °C. The supernatant contained the total chloroplast proteins. Protein concentration was determined using a PierceTM BCA protein assay kit (Thermo Fisher Scientific).

For MS analysis, equal amounts of total protein from three independent biological samples were denatured with 10 mM DTT at 56 °C for 30 min. The denatured samples were subjected to alkylation in 50 mM iodoacetamide (IAA) in the dark for 40 min. The samples were then desalted in 100 mM NH_4_HCO_3_ buffer through a Nanosep membrane (Pall Corporation, MWCO 10K). Desalted proteins were digested using trypsin (40 ng/μl trypsin and 100 mM NH_4_HCO_3_, enzyme-to-protein ratio 1:50) at 37 °C for 20 h. The cleaved peptides were then dried in a chilled CentriVap concentrator (Labconco). The peptides were resuspended in 0.1% (v/v) formic acid (FA), and subjected to nanoAcquity Ultra Performance LC (Waters) through a 20 mm trap column (C18 5 μm resin, 180 μ of 3 μl/min for 10 min, and then eluted to the analytical column (C18 1.7 μm resin, 75 μm I.D., Waters) with a flow rate of 250 nl/min under the following conditions: 140 min gradient from 8-25% of solvent B (Acetonitrile, ACN); 15 min gradient from 25-40% of solvent B; 5 min gradient from 40-90% of solvent B; 5 min washing at 90% of solvent B, and finally equilibration with 97% of solvent A for 15 min (solvent A: 0.1% FA; solvent B: 99.9% ACN/0.1% FA). After analyzing the separated peptides in a Q Exactive Mass Spectrometer (Thermo Electron Finnigan), a full MS survey scan was performed at a resolution of 70,000 at 400 m/z over the m/z range of 300-1800, with an automatic gain controls (AGC) target of 3E6 and a maximum ion injection time (IT) of 30 msec. The top 20 multiply-charged parent ions were selected under data-dependent MS/MS mode and fragmented by higher-energy collision dissociation (HCD) with a normalized collision energy of 27% and an m/z scan range of 200-2000. MS/MS detection was carried out at a resolution of 17,500 with the AGC target value of 5E5 and the maximum IT of 120 msec. Dynamic exclusion was enabled for 30 sec.

### Label-free quantitative proteomics analysis

MaxQuant software (v1.5.8.3) with an intensity-based absolute quantification (iBAQ) algorithm was used to process and analyse raw MS data as described previously^48,50,51^. Parent ion and MS2 spectra were compared against the Arabidopsis Information Resource database (http://www.arabidopsis.org/). The precursor ion tolerance was set at 7 ppm, and a fragment mass deviation of 20 ppm was allowed. The detected peptides with a minimum of six amino acids and a maximum of two missed cleavages were assigned. For both peptide and protein identification, the false discovery rate (FDR) was set to 0.01. The iBAQ intensity value was used as an accurate proxy to calculate protein amounts. Proteins detected with iBAQ intensity values in at least two out of three independent biological samples were considered meaningful.

### Statistical analyses

Numbers of biological replicates are presented in the figure legends. Statistical analyses were performed by two-tailed Student’s *t*-test or one-way analysis of variance (ANOVA) with a post-hoc Tukey’s Honest Significant Difference test. *P* values of <0.05 were considered statistically significant.

### Data availability

Data supporting the findings of this study are available within the paper and its Supplementary Information files. Source Data (gels and graphs) for Figs. 2–4 and Extended Data Figs. 3 and 4 are provided with the paper.

## Acknowledgements

We thank the Core Facility of Genomics and Bioinformatics in Shanghai Center for Plant Stress Biology (PSC) for carrying out whole-genome sequencing analysis and RNA-seq. We thank the Core Facility of Cell Biology in PSC for training students in the use of various microscopes. We also thank the Core Facility of Proteomics in PSC for conducting MS analsyis. We thank Masato Nakai (Osaka University) for providing the anti-TOC75 antibody. This research was supported by the Strategic Priority Research Program from the Chinese Academy of Sciences (Grant No. XDB27040102), the 100-Talent Program of the Chinese Academy of Sciences and the National Natural Science Foundation of China (NSFC) (Grant No. 31871397) to C.K., and by grants to H.-m.L. from the Ministry of Science and Technology (grant number MOST 109-2326-B-001-009) and Academia Sinica of Taiwan.

## Author contributions

C.K. conceived the project and designed the research: J.F., B.L., LJ.C., V.D., and S.L. conducted the experiments; J.F., B.L., V.D., P.W., HM. L., and C.K. analysed the data; C.K. wrote the manuscript with input from all authors. All authors reviewed the manuscript.

---

**Extended Data Fig. 1.**
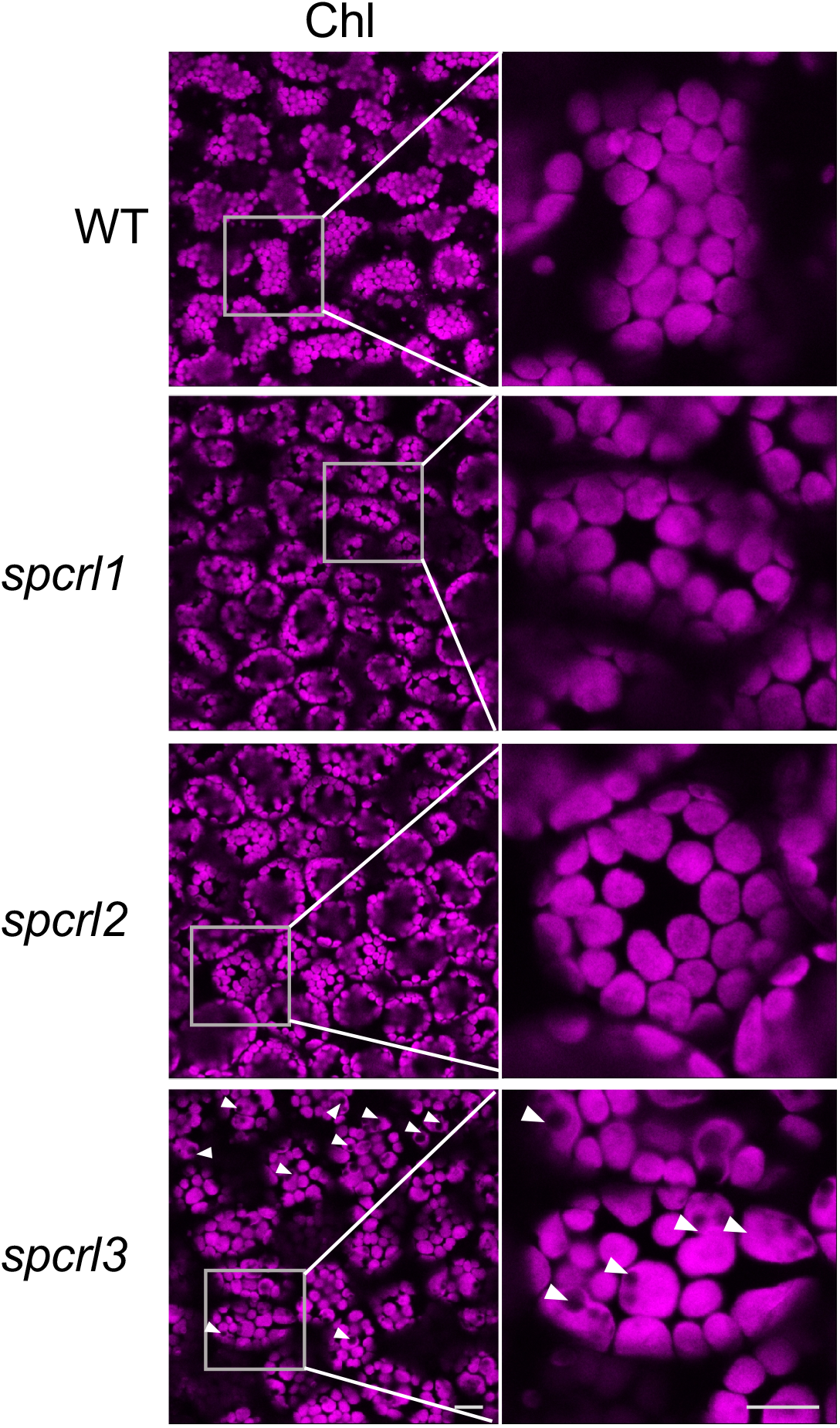
The *TIC236-6GF* mutation cannot fully rescue the *crl* mutant phenotypes. Confocal images at the same scale representing chlorophyll autofluorescence of each suppressor and WT plants (scale bar, 20 μm). The enlarged chloroplast images are shown at right (scale bar, 10 μm). White arrowheads indicate abnormally enlarged chloroplasts.

**Extended Data Fig. 2.**
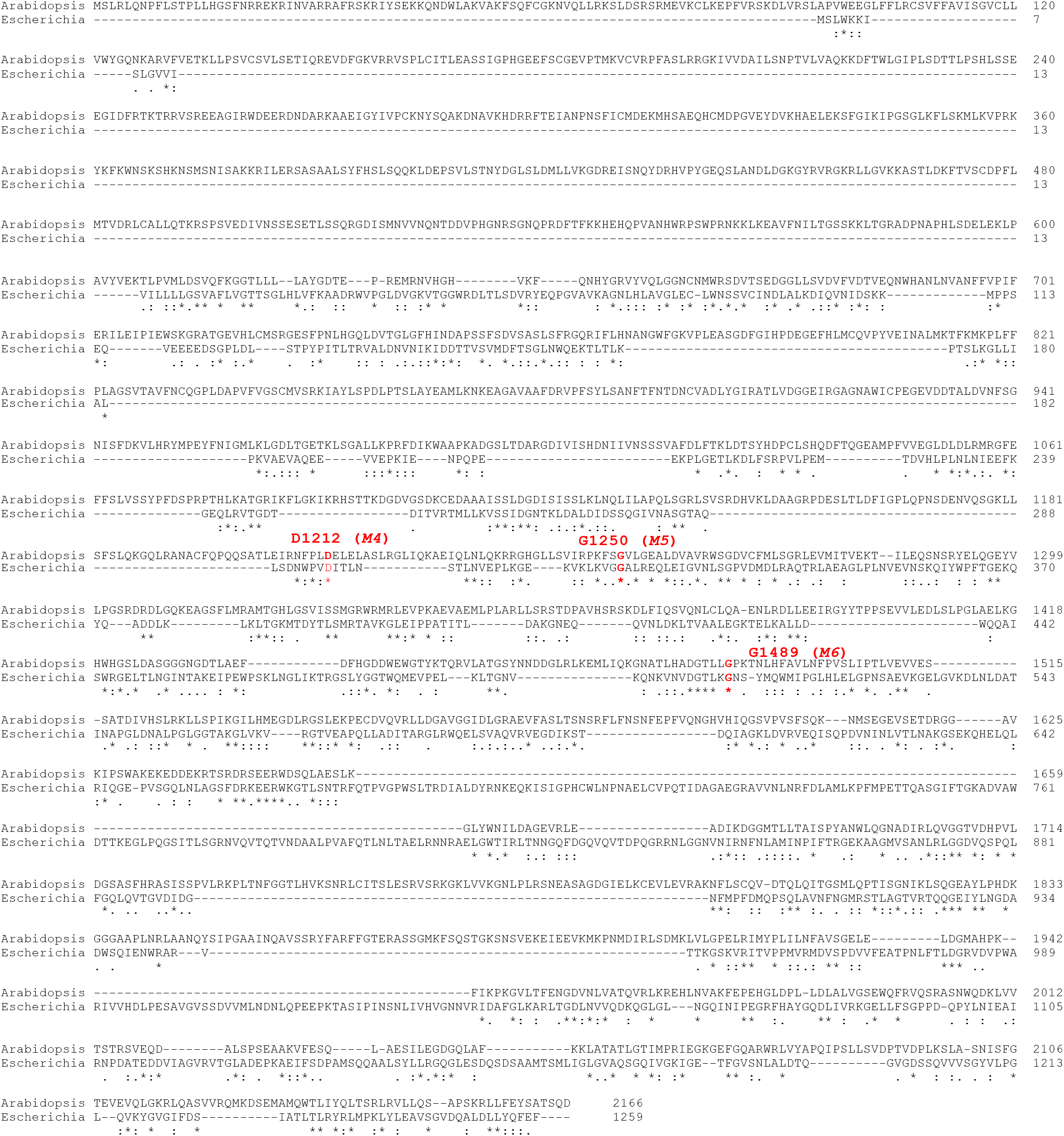
Alignment of protein sequences of Arabidopsis TIC236 and *E. coli* TamB. The amino acid sequences of TIC236 (protein accession number: NP_180137) and TamB (protein accession number: NP_418642) were obtained from NCBI (https://www.ncbi.nlm.nih.gov/protein/) and used for alignment using the Basic Local Alignment Search Tool (BLAST, https://blast.ncbi.nlm.nih.gov). Asterisks refer to conserved amino acid residues between proteins, and the mutated residues are highlighted in red. The identity of the two sequences is ∼29%.

**Extended Data Fig. 3.**
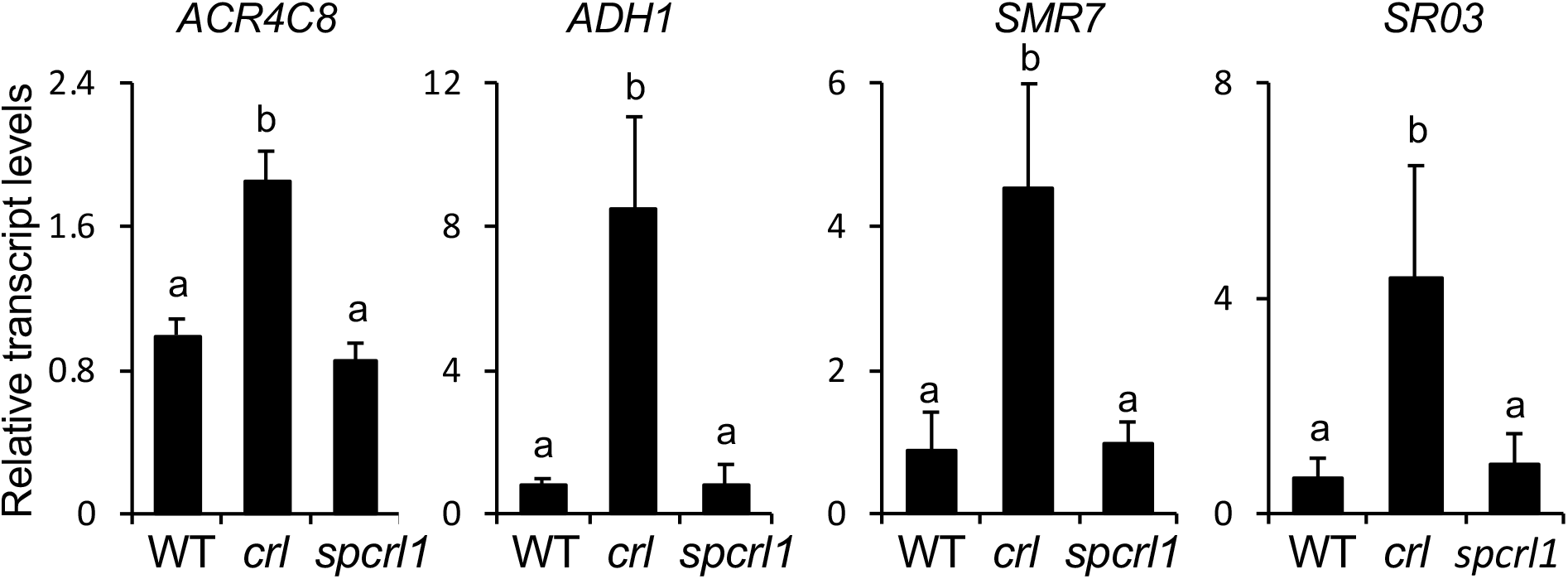
*crl*-induced stress-related nuclear genes are repressed in *spcrl1*. The relative expression levels of selected *crl*-induced genes (versus WT) were analysed in *spcrl1* mutant seedlings using qRT-PCR. These genes include *AKR4C8* (*ALDO-KETO REDUCTASE FAMILY 4 MEMBER C8*), *ADH1* (*ALCOHOL DEHYDROGENASE 1*), *SMR7* (*SIAMESE-RELATED 7*), and *SRO3* (*SIMILAR TO RCD ONE 3*). *ACTIN2* (*ACT2*) was used as an internal control. Results represent the means of three independent biological replicates and error bars indicate SD. Lower case letters indicate statistically significant differences between mean values for each genotype (P < 0.05, ANOVA with post-hoc Tukey’s Honest Significant Difference test).

**Extended Data Fig. 4.**
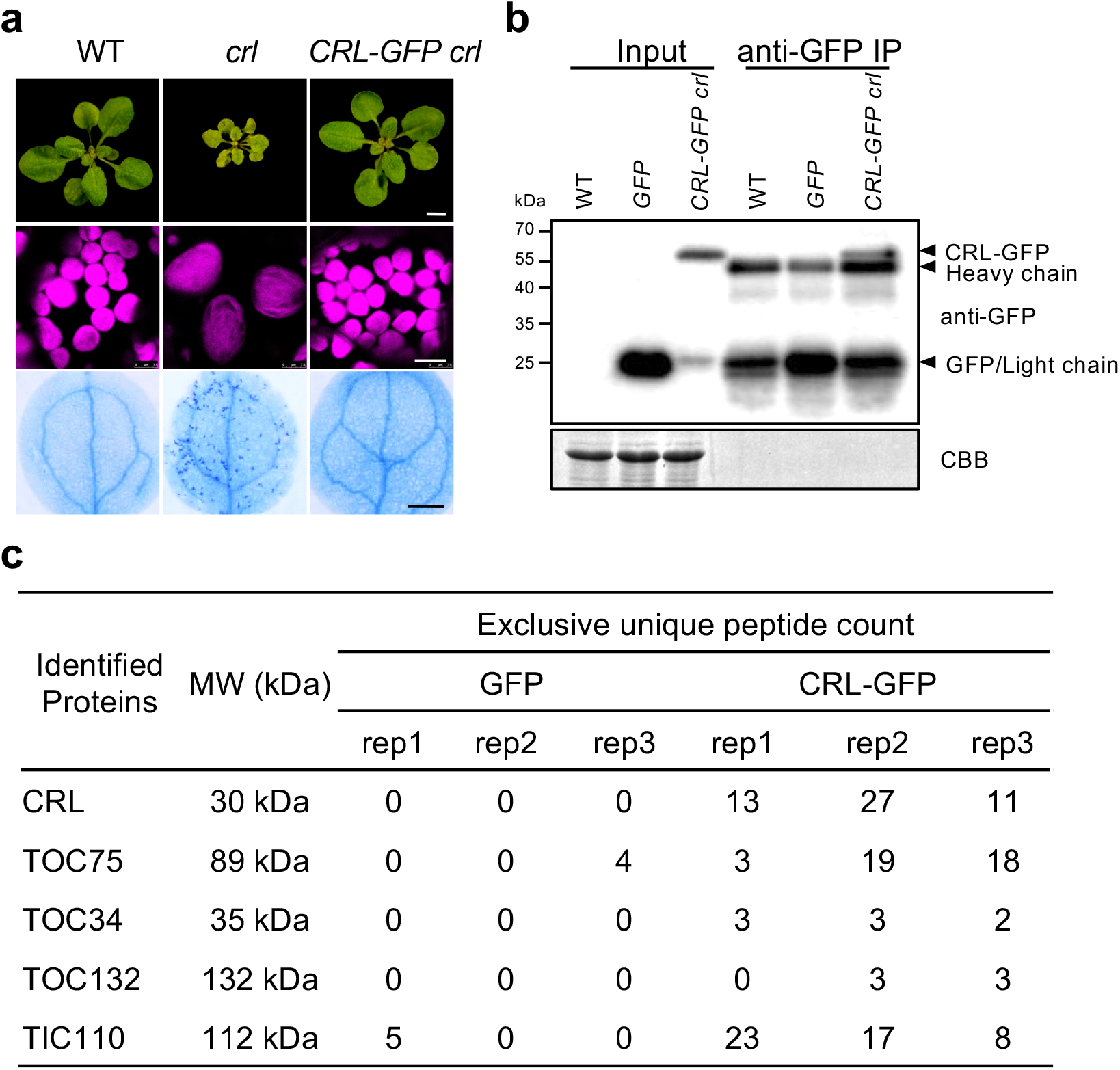
The biologically active CRL-GFP fusion protein unveils CRL-associated proteins. **a**, Top: images represent 21-d-old WT, *crl*, and *35S:CRL- GFP crl* (*CRL-GFP crl*) plants. Scale bar, 5 mm. Middle: confocal images of chlorophyll autofluorescence of 5-d-old cotyledons (scale bar, 10 µm). Bottom: localized cell death in 10-d-old cotyledons, as visualized by TB staining (scale bar, 0.5 mm). **b**, Western blot of Co-IP using 14-d-old WT, *35S:GFP* (*GFP*), and *CRL- GFP crl* plants. GFP-conjugated Dynabeads were used to pull down CRL-GFP and its associated proteins. The proteins were subjected to MS analysis after digestion. In parallel, equal amounts of proteins were used for Western blot analysis. Heavy and light chains of the GFP antibody are indicated. Equal protein loading is shown by CBB staining. **c**, List of proteins showing TOC and TIC components, together with CRL, which were detected at least twice in the eluates from *CRL-GFP crl* but not in *GFP* samples.

**Extended Data Fig. 5.**
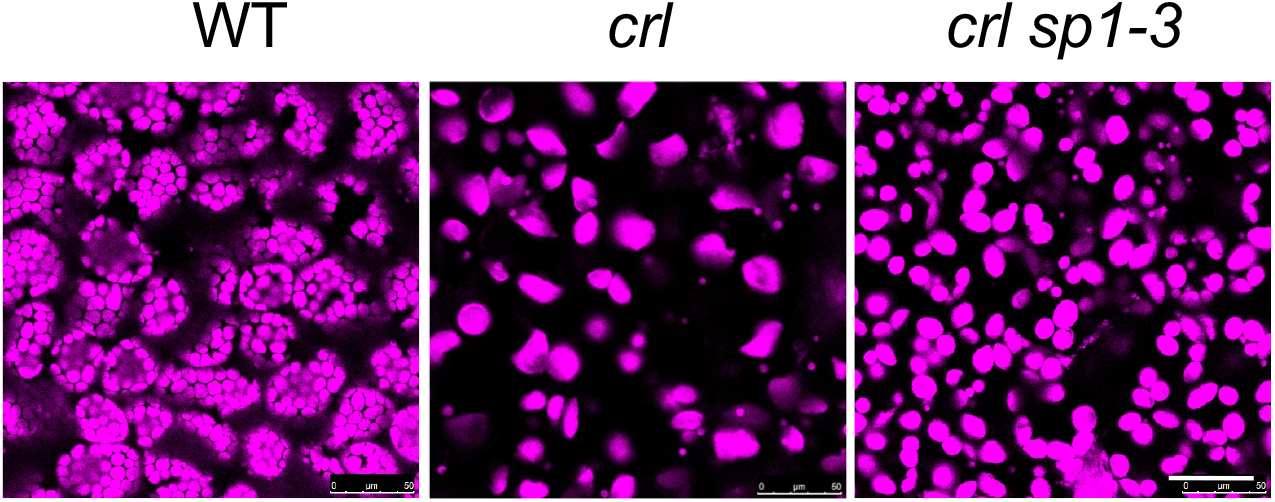
*sp1-3* significantly restores plastid division in *crl*. Same-scale confocal images representing the chlorophyll autofluorescence in mesophyll cells of 5-d-old seedlings (scale bar, 50 μm).

**Extended Data Fig. 6.**
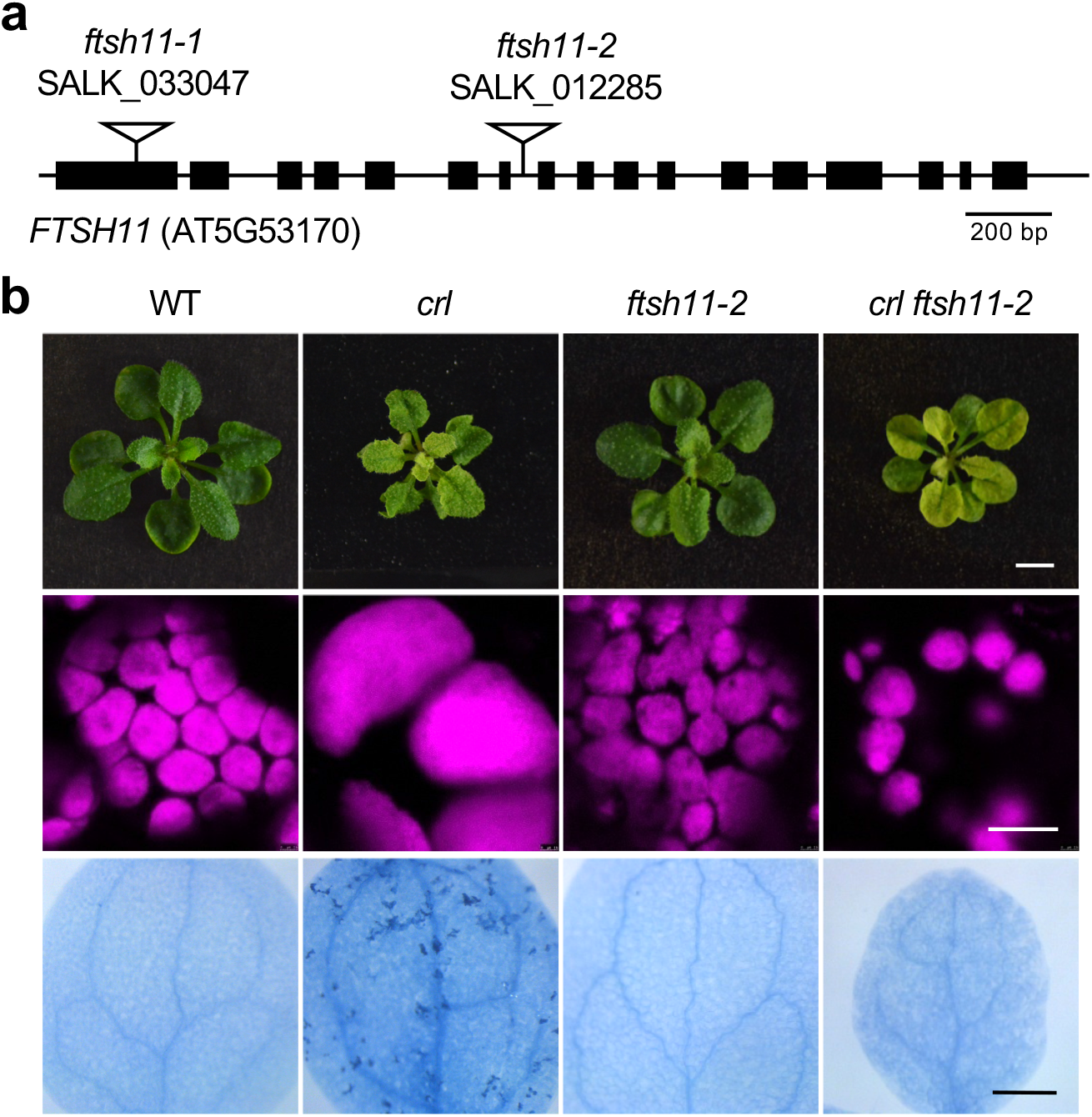
*ftsh11-2* significantly restores plastid division in *crl.* **a**, Schematic representation of the FTSH11 gene (accession number: AT5G53170). Exons and introns are shown as black boxes and black lines between exons, respectively. The inverted triangles indicate the T-DNA insertion sites of the *ftsh11-1* (SALK_033047) and *ftsh11-2* (SALK_012285) mutants. **b**, Top: representative plant images of 21-d-old WT, *crl*, *ftsh11-2*, and *crl ftsh11-2* are shown (top panel). Middle: confocal images of chlorophyll signals from mesophyll cells (scale bar, 10 µm). Bottom: cell death in cotyledons, as visualized via TB staining (scale bar, 0.5 mm).

**Extended Data Fig. 7.**
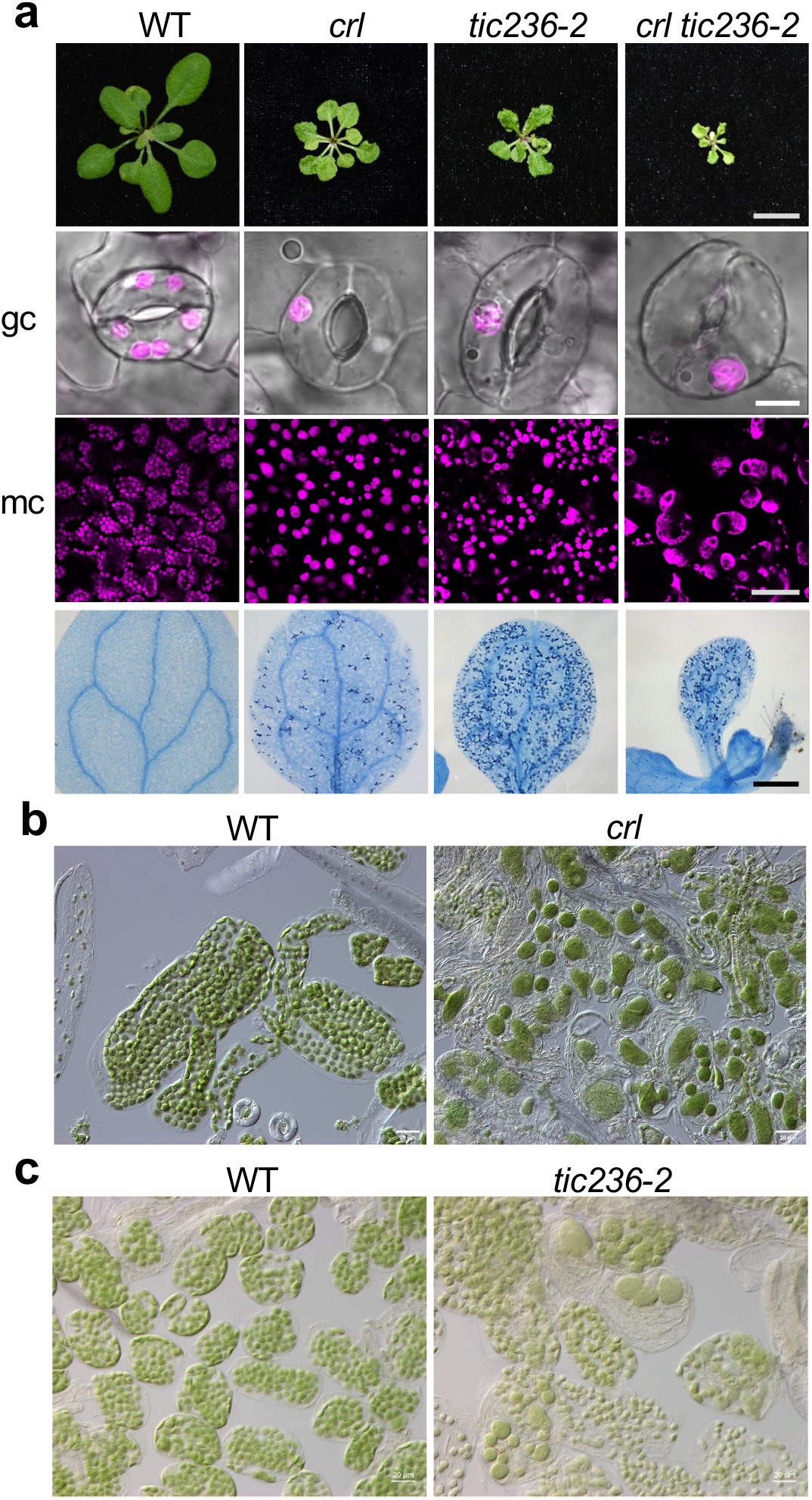
The *tic236*-*2* mutant, like the *crl* mutant, exhibits cell lesion and defective plastid division phenotypes. **a**, Top: representative images of the rosettes of 21-d-old plants (scale bars, 1 cm). Middle two panels: confocal images of chlorophyll autofluorescence and the cognate bright-field in 5-d-old cotyledons for guard cells (gc) and mesophyll cells (mc) (scale bars, 5 μm and 40 μm for gc and mc, respectively). Bottom: cell death in 10-d-old cotyledons, as visualized by TB staining (scale bar, 0.5 mm). **b** and **c**, Mesophyll cells of mature *crl* (**b**) and *tic236-2* (**c**) plants were observed by differential interference contrast (DIC) optics (scale bar: 20 μm).

**Extended Data Fig. 8.**
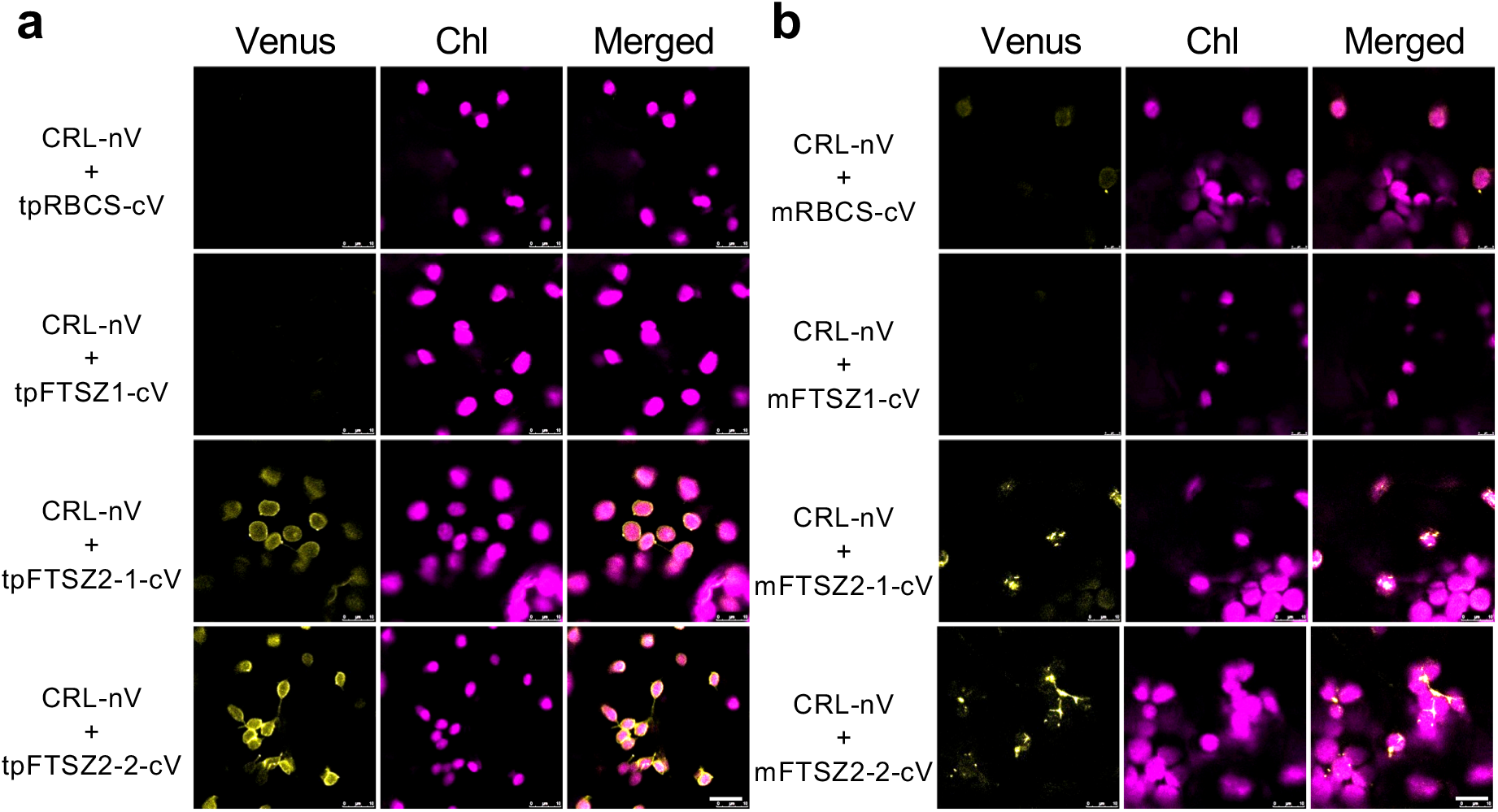
CRL interacts with transit peptides of FTSZ proteins. **a** and **b**, Split-Venus constructs of CRL and the transit peptide (tp) lacking mature protein region (**a**) or tp-deleted mature protein (m) (**b**) of either RBCS, FTSZ1, FTSZ1, FTSZ2-1, or FTSZ2-2 were transiently coexpressed in *N. benthamiana* leaves. Fluorescence signal of the integrated Venus protein was monitored by confocal microscopy. Scale bar, 10 μm.

## Main References

1 Schwenkert, S., Dittmer, S. & Soll, J. Structural components involved in plastid protein import. Essays Biochem 62, 65–75, doi:10.1042/EBC20170093 (2018).

2 Richardson, L. G. L. & Schnell, D. J. Origins, function, and regulation of the TOC-TIC general protein import machinery of plastids. J Exp Bot 71, 1226–1238, doi:10.1093/jxb/erz517 (2020).

3 Chen, Y. L. et al. TIC236 links the outer and inner membrane translocons of the chloroplast. Nature 564, 125–129, doi:10.1038/s41586-018-0713-y (2018).

4 Patel, R., Hsu, S. C., Bedard, J., Inoue, K. & Jarvis, P. The Omp85-related chloroplast outer envelope protein OEP80 is essential for viability in Arabidopsis. Plant Physiol 148, 235–245, doi:10.1104/pp.108.122754 (2008).

5 Selkrig, J. et al. Discovery of an archetypal protein transport system in bacterial outer membranes. Nat Struct Mol Biol 19, 506–510, S501, doi:10.1038/nsmb.2261 (2012).

6 Selkrig, J. et al. Conserved features in TamA enable interaction with TamB to drive the activity of the translocation and assembly module. Sci Rep 5, 12905, doi:10.1038/srep12905 (2015).

7 Josts, I. et al. The structure of a conserved domain of TamB reveals a hydrophobic beta taco fold. Structure 25, 1898–1906 e1895, doi:10.1016/j.str.2017.10.002 (2017).

8 Schnell, D. J. Exit route evolved into entry path in plants. Nature 564, 45–46, doi:10.1038/d41586-018-07426-6 (2018).

9 Osteryoung, K. W. & Pyke, K. A. Division and dynamic morphology of plastids. Annu Rev Plant Biol 65, 443–472, doi:10.1146/annurev-arplant-050213-035748 (2014).

10 Simkova, K. et al. The chloroplast division mutant caa33 of Arabidopsis thaliana reveals the crucial impact of chloroplast homeostasis on stress acclimation and retrograde plastid-to-nucleus signaling. Plant J 69, 701–712, doi:10.1111/j.1365-313X.2011.04825.x (2012).

11 Bruggeman, Q., Raynaud, C., Benhamed, M. & Delarue, M. To die or not to die? Lessons from lesion mimic mutants. Front Plant Sci 6, 24, doi:10.3389/fpls.2015.00024 (2015).

12 Lorrain, S., Vailleau, F., Balague, C. & Roby, D. Lesion mimic mutants: keys for deciphering cell death and defense pathways in plants? Trends Plant Sci 8, 263–271, doi:10.1016/S1360-1385(03)00108-0 (2003).

13 Li, B. et al. FATTY ACID DESATURASE5 is required to induce autoimmune responses in gigantic chloroplast mutants of Arabidopsis. Plant Cell 32, 3240–3255, doi:10.1105/tpc.20.00016 (2020).

14 Hudik, E. et al. Chloroplast dysfunction causes multiple defects in cell cycle progression in the Arabidopsis crumpled leaf mutant. Plant Physiol 166, 152–167, doi:10.1104/pp.114.242628 (2014).

15 Asano, T. et al. A mutation of the CRUMPLED LEAF gene that encodes a protein localized in the outer envelope membrane of plastids affects the pattern of cell division, cell differentiation, and plastid division in Arabidopsis. Plant J 38, 448–459, doi:10.1111/j.1365-313X.2004.02057.x (2004).

16 Grossman, A. R., Schaefer, M. R., Chiang, G. G. & Collier, J. L. The phycobilisome, a light-harvesting complex responsive to environmental conditions. Microbiol Rev 57, 725–749 (1993).

17 Wang, F. et al. The Arabidopsis CRUMPLED LEAF protein, a homolog of the cyanobacterial bilin lyase, retains the bilin-binding pocket for a yet unknown function. Plant J 104, 964–978, doi:10.1111/tpj.14974 (2020).

18 Ling, Q. et al. Ubiquitin-dependent chloroplast-associated protein degradation in plants. Science 363, 836-+, doi:10.1126/science.aav4467 (2019).

19 Shanmugabalaji, V. & Kessler, F. CHLORAD: Eradicating translocon components from the outer membrane of the chloroplast. Mol Plant 12, 467–469, doi:10.1016/j.molp.2019.03.002 (2019).

20 Ling, Q., Huang, W., Baldwin, A. & Jarvis, P. Chloroplast biogenesis is regulated by direct action of the ubiquitin-proteasome system. Science 338, 655–659, doi:10.1126/science.1225053 (2012).

21 Ling, Q. & Jarvis, P. Regulation of chloroplast protein import by the ubiquitin E3 ligase SP1 is important for stress tolerance in plants. Curr Biol 25, 2527–2534, doi:10.1016/j.cub.2015.08.015 (2015).

22 Adam, Z. et al. The chloroplast envelope protease FTSH11 - Interaction with CPN60 and identification of potential substrates. Front Plant Sci 10, 428, doi:10.3389/fpls.2019.00428 (2019).

23 Suzuki, K. et al. Plastid chaperonin proteins Cpn60 alpha and Cpn60 beta are required for plastid division in Arabidopsis thaliana. BMC Plant Biol 9, 38, doi:10.1186/1471-2229-9-38 (2009).

24 Chen, J., Xin, Z. & Burke, J. The conserved role of FtsH11 protease in protection of photosynthetic system from high temperature stress in higher plants. Photosynthesis Res 91, 308–308, doi: 10.3389/fpls.2019.00428 (2007).

25 Chen, J., Burke, J. J. & Xin, Z. Chlorophyll fluorescence analysis revealed essential roles of FtsH11 protease in regulation of the adaptive responses of photosynthetic systems to high temperature. BMC Plant Biol 18, 11, doi:10.1186/s12870-018-1228-2 (2018).

26 Chen, Y. et al. Plant cells without detectable plastids are generated in the crumpled leaf mutant of Arabidopsis thaliana. Plant Cell Physiol 50, 956–969, doi:10.1093/pcp/pcp047 (2009).

27 Kessler, F. & Blobel, G. Interaction of the protein import and folding machineries of the chloroplast. Proc Natl Acad Sci U S A 93, 7684–7689, doi:10.1073/pnas.93.15.7684 (1996).

28 Klasek, L., Inoue, K. & Theg, S. M. Chloroplast chaperonin-mediated targeting of a thylakoid membrane protein. Plant Cell 32, 3884–3901, doi:10.1105/tpc.20.00309 (2020).

29 Matsushima, R. et al. Amyloplast-localized SUBSTANDARD STARCH GRAIN4 protein influences the size of starch grains in rice endosperm. Plant Physiol 164, 623–636, doi:10.1104/pp.113.229591 (2014).

30 Zhang, J. et al. Maize defective kernel5 is a bacterial TamB homologue required for chloroplast envelope biogenesis. J Cell Biol 218, 2638–2658, doi:10.1083/jcb.201807166 (2019).

## Method References

31 Kim, Y., Schumaker, K. S. & Zhu, J. K. EMS mutagenesis of Arabidopsis. Methods Mol Biol 323, 101–103, doi:10.1385/1-59745-003-0:101 (2006).

32 Cox, M. P., Peterson, D. A. & Biggs, P. J. SolexaQA: At-a-glance quality assessment of Illumina second-generation sequencing data. BMC Bioinformatics 11, 485, doi:10.1186/1471-2105-11-485 (2010).

33 Li, H. & Durbin, R. Fast and accurate short read alignment with Burrows-Wheeler transform. Bioinformatics 25, 1754–1760, doi:10.1093/bioinformatics/btp324 (2009).

34 Li, H. et al. The Sequence Alignment/Map format and SAMtools. Bioinformatics 25, 2078–2079, doi:10.1093/bioinformatics/btp352 (2009).

35 Danecek, P. et al. The variant call format and VCFtools. Bioinformatics 27, 2156–2158, doi:10.1093/bioinformatics/btr330 (2011).

36 Schneeberger, K. et al. SHOREmap: simultaneous mapping and mutation identification by deep sequencing. Nat Methods 6, 550–551, doi:10.1038/nmeth0809-550 (2009).

37 Sun, H. & Schneeberger, K. SHOREmap v3.0: fast and accurate identification of causal mutations from forward genetic screens. Methods in molecular biology (Clifton, N.J.) 1284, 381–395, doi:10.1007/978-1-4939-2444-8_19 (2015).

38 Wang, L. et al. Singlet oxygen- and EXECUTER1-mediated signaling is initiated in grana margins and depends on the protease FtsH2. Proc Natl Acad Sci U S A 113, E3792–3800, doi:10.1073/pnas.1603562113 (2016).

39 Lee, D. W. et al. Molecular Mechanism of the Specificity of Protein Import into Chloroplasts and Mitochondria in Plant Cells. Molecular plant 12, 951–966, doi:10.1016/j.molp.2019.03.003 (2019).

40 Emanuelsson, O., Nielsen, H. & von Heijne, G. ChloroP, a neural network-based method for predicting chloroplast transit peptides and their cleavage sites. Protein Sci 8, 978–984 (1999).

41 Schmitz, A. J., Glynn, J. M., Olson, B. J., Stokes, K. D. & Osteryoung, K. W. Arabidopsis FtsZ2-1 and FtsZ2-2 are functionally redundant, but FtsZ-based plastid division is not essential for chloroplast partitioning or plant growth and development. Mol Plant 2, 1211–1222, doi:10.1093/mp/ssp077 (2009).

42 Chu, C. C. & Li, H. m. Determining the location of an Arabidopsis chloroplast protein using in vitro import followed by fractionation and alkaline extraction. Methods Mol Biol 774, 339–350, doi:10.1007/978-1-61779-234-2_20 (2011).

43 Pyke, K. A. & Leech, R. M. Rapid Image Analysis Screening Procedure for Identifying Chloroplast Number Mutants in Mesophyll Cells of Arabidopsis thaliana (L.) Heynh. Plant physiology 96, 1193–1195, doi:10.1104/pp.96.4.1193 (1991).

44 Dogra, V. et al. FtsH2-Dependent Proteolysis of EXECUTER1 Is Essential in Mediating Singlet Oxygen-Triggered Retrograde Signaling in Arabidopsis thaliana. Front Plant Sci 8, 1145, doi:10.3389/fpls.2017.01145 (2017).

45 Trapnell, C., Pachter, L. & Salzberg, S. L. TopHat: discovering splice junctions with RNA-Seq. Bioinformatics 25, 1105–1111, doi:10.1093/bioinformatics/btp120 (2009).

46 Robinson, M. D., McCarthy, D. J. & Smyth, G. K. edgeR: a Bioconductor package for differential expression analysis of digital gene expression data. Bioinformatics 26, 139–140, doi:10.1093/bioinformatics/btp616 (2010).

47 Livak, K. J. & Schmittgen, T. D. Analysis of relative gene expression data using real-time quantitative PCR and the 2(-Delta Delta C(T)) Method. Methods 25, 402–408, doi:10.1006/meth.2001.1262 (2001).

48 Dogra, V., Duan, J., Lee, K. P. & Kim, C. Impaired PSII proteostasis triggers a UPR-like response in the var2 mutant of Arabidopsis. J Exp Bot 70, 3075–3088, doi:10.1093/jxb/erz151 (2019).

49 Kauss, D., Bischof, S., Steiner, S., Apel, K. & Meskauskiene, R. FLU, a negative feedback regulator of tetrapyrrole biosynthesis, is physically linked to the final steps of the Mg(++)-branch of this pathway. FEBS Lett 586, 211–216, doi:10.1016/j.febslet.2011.12.029 (2012).

50 Luber, C. A. et al. Quantitative proteomics reveals subset-specific viral recognition in dendritic cells. Immunity 32, 279–289, doi:10.1016/j.immuni.2010.01.013 (2010).

51 Schwanhausser, B. et al. Global quantification of mammalian gene expression control. Nature 473, 337–342, doi:10.1038/nature10098 (2011).

